# Genome-wide association studies of brain structure and function in the UK Biobank

**DOI:** 10.1101/178806

**Authors:** Lloyd T. Elliott, Kevin Sharp, Fidel Alfaro-Almagro, Sinan Shi, Karla Miller, Gwenaëlle Douaud, Jonathan Marchini, Stephen Smith

## Abstract

The genetic basis of brain structure and function is largely unknown. We carried out genome-wide association studies of 3,144 distinct functional and structural brain imaging derived phenotypes in UK Biobank (discovery dataset 8,428 subjects). We show that many of these phenotypes are heritable. We identify 148 clusters of SNP-imaging associations with lead SNPs that replicate at p<0.05, when we would expect 21 to replicate by chance. Notable significant and interpretable associations include: iron transport and storage genes, related to changes in T2* in subcortical regions; extracellular matrix and the epidermal growth factor genes, associated with white matter micro-structure and lesion volume; genes regulating mid-line axon guidance development associated with pontine crossing tract organisation; and overall 17 genes involved in development, pathway signalling and plasticity. Our results provide new insight into the genetic architecture of the brain with relevance to complex neurological and psychiatric disorders, as well as brain development and aging. The full set of results is available on the interactive Oxford Brain Imaging Genetics (BIG) web browser.

## Main text

Brain structure and function are known to vary between individuals in the human population and can be measured non-invasively using Magnetic Resonance Imaging (MRI). Disease effects seen in MRI data have been identified in many neurological and psychiatric disorders such as Alzheimer’s disease, Parkinson’s disease, schizophrenia, bipolar disorder and autism^1^. MRI can provide intermediate or endo-phenotypes that can be used to assess the genetic architecture of such disorders.

Structural MRI measures of brain anatomy include tissue and structure volumes, such as total grey matter volume and hippocampal volume, while other MRI modalities allow the mapping of different biological markers such as venous vasculature, microbleeds and aspects of white matter (WM) micro-structure. Brain function is typically measured using task-based functional MRI (tfMRI) in which subjects perform tasks or experience sensory stimuli, and uses imaging sensitive to local changes in blood oxygenation and flow caused by brain activity in grey matter. Brain connectivity can be divided into functional connectivity, where spontaneous temporal synchronisations between brain regions are measured using fMRI with subjects scanned at rest, and structural connectivity, measured using diffusion MRI (dMRI), which images the physical connections between brain regions based on how water molecules diffuse within white matter tracts. For those not familiar with the neuroimaging field, we have provided a glossary in **Supplementary Note 1**.

A new resource for relating neuroimaging measures to genetics is UK Biobank, a rich, long-term prospective epidemiological study of 500,000 volunteers^2^. Participants were 40-69 years of age at baseline recruitment, a major aim being to acquire as rich data as possible before disease onset. Identification of disease risk factors and early markers will increase over time with emerging clinical outcomes^3^. A brain and body imaging extension will scan 100,000 participants by 2020, with brain imaging including three structural modalities, resting and task fMRI, and diffusion MRI^4^ (**Supplementary Table 1**). A fully automated image processing pipeline removes artefacts and renders images comparable across modalities and participants. The pipeline also generates thousands of image-derived phenotypes (IDPs), distinct individual measures of brain structure and function ^5^ Example IDPs include the volume of grey matter in many distinct brain regions, and measures of functional and structural connectivity between specific pairs of brain areas. The combination of very large subject numbers with richly multimodal imaging data collected on homogeneous imaging hardware and software is a unique feature of UK Biobank.

Another key component of the UK Biobank resource has been the collection of genome-wide genetic data using a purpose-designed genotyping array. A custom quality control, phasing and imputation pipeline was developed to address the challenges specific to the experimental design, scale, and diversity of the UK Biobank dataset. The genetic data was publicly released in July 2017 and consists of ~96 million genetic variants in ~500,000 participants.^6^

Joint analysis of the genetic and brain imaging datasets produced by UK Biobank presents a unique opportunity for uncovering the genetic bases of brain structure and function, including genetic factors relating to brain development, aging and disease. In this study, we carried out genome-wide association studies (GWAS) for 3,144 IDPs, covering the entire brain and including “multi-modal” information of grey matter volume, area and thickness, white matter connections and functional connectivity, at 11,734,353 SNPs (single-nucleotide polymorphisms) in up to 8,428 individuals having both genetic and brain imaging data. We used two separate sets of data from UK Biobank to evaluate replication of significant genetic associations from the discovery phase. We also carried out multi-trait GWAS, SNP-heritability analysis, genetic correlation analysis of IDPs with brain-related traits and an analysis of enrichment of genomic regions with different functions. Previous large-scale GWAS imaging studies have focussed on narrower ranges of phenotypes including studies of: grey matter volume in 7 localised regions of the subcortical brain by combining data across >50 different studies^7,8^ whole brain grey matter volumes and thicknesses by combining data from 59 acquisition sites ^9^; and cortico-cortical white matter connections in healthy young adult twins^10^. We expect that homogeneous image aquistion and genetic data assay in UK Biobank will have a positive impact on the power of our study.

The full set of results are available on the Oxford Brain Imaging Genetics (BIG) web browser that allows users to browse associations by SNP, gene or phenotype. This browser was built from the PheWeb code base and extended to allow easier searching of phenotypes. In addition to the brain IDP GWAS results, the browser also includes GWAS results from more than 2,500 other traits and diseases (see **URLs**).

### Heritability and genetic correlations of IDPs

**Figure 1** shows the estimated SNP-heritability (*h*^*2*^) of all IDPs and whether *h*^*2*^ is significantly different from 0 at the nominal 5% significance level (see also **Supplementary Table 2 and Supplementary Figure 1**). 1,578 of 3,144 IDPs show significant SNP-heritability. Of the structural MRI IDPs, volumetric measures are the most heritable and cortical thicknesses the least. Of the diffusion MRI measures, the tractography-based IDPs show lower heritability than the tract-skeleton-based IDPs. The resting-state fMRI functional connectivity edges show the lowest levels of SNP-heritability, with just 235 of 1,771 IDPs significant, which is consistent with additive heritability estimates from twin studies of network edges from fMRI and MEG in the 11 Human Connectome Project ^11^. However, 4 of the 6 rfMRI ICA features (estimated as data-driven reductions of this full set of fMRI edges) are much more highly heritable. In contrast the resting-state node amplitude IDPs do mostly show significant evidence of SNP-heritability; the task fMRI IDPs do not.

**Figure 1:**
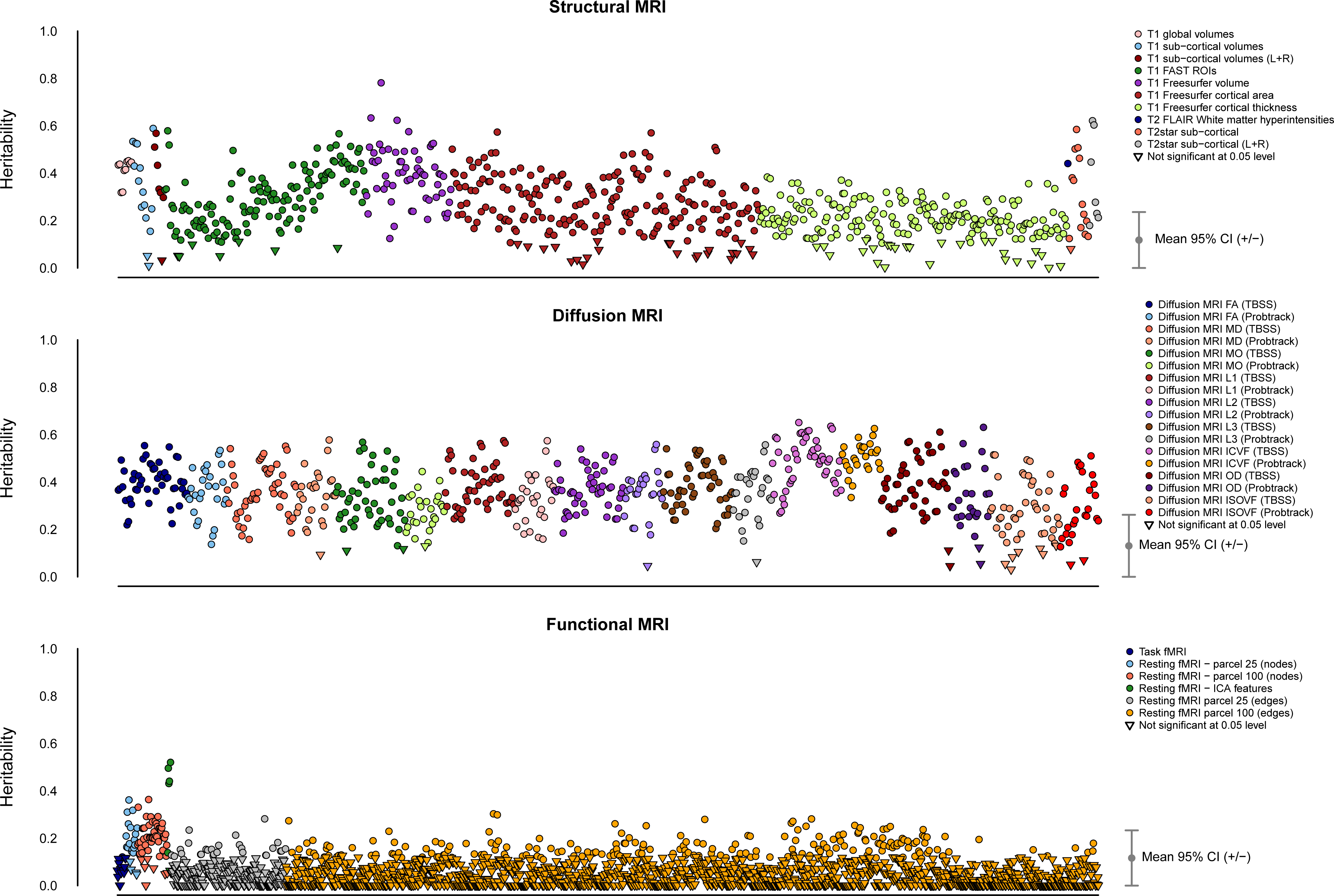
Estimated heritability of IDPs. Estimated heritability (y-axis) of all of the IDPs analyzed. IDPs have been split into three broad groups: Structural MRI (top), Diffusion MRI (middle) and Functional MRI (bottom). Points are colored according to IDP groups. Circles and inverted triangles are used to identify IDPs that do/do not have heritability significantly different from 0 at the 5% significance level. The mean 95% confidence interval (CI) is also indicated to the right of each group of IDPs.

We found lower levels of SNP-heritability for sub-cortical volumes than previously 12 14 estimated in twin studies ^12–14^ (**Supplementary Figure 2**). This is typical of many traits in the literature^15^ and maybe due to twin study estimates being upwardly biased 16 17 due to gene-gene and gene-environment interactions^16,17^, or downward bias of SNP-heritability due to uncaptured rare genetic variation. We also compared the GWAS results for 7 subcortical volumes with those obtained by the ENIGMA consortium, via a genetic correlation analysis (**Supplementary Table 3**). We find a strong correlation between the studies, suggesting no major differences between how these phenotypes have been measured. In all cases a perfect genetic correlation of 1 lies within the 95% confidence intervals.

**Supplementary Figure 3** shows the genetic correlations, together with the raw phenotype correlations, for several groups of analysed IDPs. These plots show that there is a range of both strong and weak, positive and negative genetic correlations between the IDPs.

### Significant associations between IDPs and SNPs

In all analyses we estimated genetic effects with respect to the number of copies of the *non-reference allele.* Using a minor allele frequency filter of 1% and a -log_10_ p-value threshold of 7.5, we found 1,262 significant associations between SNPs and the IDPs. These associations span all classes of IDPs, except task fMRI (**Supplementary Table 4**), with the swMRI T2* group showing a relatively large number of associations. The -log10 p-value threshold of 7.5 controls for the number of tests carried out across SNPs and takes into account the correlation structure between genetic variants. 844 and 455 of these 1,262 associations replicated at the 5% significance level using our two smaller replication datasets (**Methods** and **Supplementary Table 5**). Some associated genetic loci overlap across IDPs; we estimate that there are approximately 427 distinct associated genetic regions (“clusters”), and 148 of these “clusters” have a lead SNP that replicates at the 5% level in our replication set of 3,456 participants, and 91 below a 5% False Discovery Rate (FDR) threshold. We would expect ~21 of the lead SNPs in the 148 clusters to replicate under a null hypothesis of no association.

At a threshold of -log10 p-value > 11, which additionally corrects for all 3,144 GWAS carried out (see **Methods**), we find 368 significant associations between genetic regions and distinct IDPs (**Supplementary Table 6, Supplementary Figure 4**). These associations with 78 unique SNPs can be grouped together into 38 distinct clusters by grouping across IDPs (**Table 1**). Taking our lead SNP in each of the 38 regions, we find that all 38 have p<0.05 in our replication set of 3,456 participants, and all 38 are significant at 5% FDR. We found no appreciable change in these GWAS results when we included a set of potential body confound measures in addition to the main set of imaging confound measures (see **Methods** and **Supplementary Figure 5**). We also carried out a Winner’s Curse corrected post-hoc power analysis that agrees well with the results of our replication studies. (**Supplementary Note 2**).

**Table 1:** Summary of most highly associated SNP-IDP clusters. The table summarises the 38 clusters of SNP-IDP associations. For each cluster the most significant association between a SNP and an IDP is detailed by the chromosome, rsID, base-pair position, SNP alleles, non-reference allele frequency, p-value in the discovery sample and the replication p-values. The locus column details a gene if the SNP is in that gene. If we found a coding SNP or eQTL in high LD with the lead SNP, then this is reported instead.

| cluster index | cluster name | # IDPs | top IDP | chr | RSID | position | locus | ref allele | nonref allele | nonref AF | p value | replication p-value (N=3456) | replication p-value (N=930) | GTEx eQTL |
| --- | --- | --- | --- | --- | --- | --- | --- | --- | --- | --- | --- | --- | --- | --- |
| 1 | Volume Cerebellum VIIIa (vermis) | 1 | T1_FAST_ROIs_V_cerebellum_VIIIa | 1 | rs76934732 | 76013268 | SLC44A5 | G | A | 0.145 | 8.51E-13 | 6.10E-04 | 5.22E-02 | SLC44A5 ACADM |
| 2 | dMRI Corpus callosum (genu) | 1 | dMRI_TBSS_ICVF_Genu_of_corpus_callosum | 1 | rs2365715 | 156615114 | BCAN | A | G | 0.388 | 5.38E-12 | 4.50E-03 | 1.33E-02 | BCAN. APOA1BP, SYT11 |
| 3 | Volume WM lesions | 1 | T2_FLAIR_BIANCA_WMH_volume | 2 | rs3762515 (5' UTR) | 56150864 | EFEMP1 | C | T | 0.0959 | 4.27E-13 | 1.18E-02 | 4.84E-01 |  |
| 4 | rfMRI Cortical and cerebellar motor nodes and edges | 2 | NODEamps25_0012 | 2 | rs60873293 | 114092549 | intergenic | G | T | 0.217 | 9.86E-15 | 3.10E-07 | 9.50E-02 | AC016745.3, RP11-480C16.1 |
| 5 | T2* Pallidum | 1 | SWI_T2*_pallidum_L+R | 2 | rs6740926 | 190326498 | WDR75 | C | T | 0.038 | 1.31E-14 | 3.50E-09 | 3.78E-04 | WDR75 |
| 6 | rfMRI Middle temporal sulcus nodes and edges | 2 | netmat_ICA_003 | 3 | rs35124509 (missense) | 89521693 | EPHA3 | T | C | 0.3853 | 4.49E-22 | 3.27E-09 | 3.73E-03 | EPHA3 |
| 7 | T2* Putamen and pallidum | 6 | SWI_T2*_putamen_L+R | 3 | rs4428180 | 133466374 | TF | A | G | 0.152 | 2.23E-22 | 6.11E-07 | 1.03E-03 | TF |
| 8 | rfMRI Prefrontal and parietal edges | 1 | netmat_ICA_002 | 3 | rs2279829 (3' UTR) | 147106319 | ZIC4 | C | T | 0.221 | 8.34E-12 | 5.46E-05 | 2.51E-03 |  |
| 9 | dMRI Superior cerebellar peduncles | 8 | dMRI_TBSS_ICVF_Superior_cerebellar_peduncle_L | 4 | rs4697414 | 23724255 | RP11-380P13.2 | C | T | 0.823 | 5.83E-24 | 1.33E-06 | 4.63E-02 | RP13-497K6.1, RP11-380P13.2 |
| 10 | Volume Putamen, ventral striatum, cerebellum VIIIb, IX, X; T2* Pallidum; dMRI Cerebral peduncles | 20 | IDP_T1_FAST_ROIs_L_ventral_striatum | 4 | rs13107325 (missense) | 103188709 | SLC39A8 | C | T | 0.073 | 1.04E-42 | 6.64E-20 | 8.97E-06 |  |
| 11 | dMRI Most WM tracts | 199 | dMRI_ProbtrackX_ICVF_illf_r | 5 | rs67827860 | 82860485 | VCAN | C | T | 0.188 | 4.06E-37 | 3.93E-12 | 2.19E-04 |  |
| 12 | rfMRI Parietal and prefrontal edges | 1 | netmat_ICA_004 | 5 | rs7442779 | 92788278 | NR2F1-AS1 | A | G | 0.05 | 8.18E-15 | 1.90E-04 | 4.04E-02 |  |
| 13 | dMRI Corpus callosum (genu, body, splenium) | 7 | dMRI_TBSS_ICVF_Genu_of_corpus_callosum | 5 | rs4150221 | 139719991 | HBEGF | T | C | 0.264 | 8.43E-20 | 1.72E-09 | 4.06E-02 | SRA1 |
| 14 | T2* Putamen | 3 | SWI_T2*_putamen_L+R | 6 | rs1800562 (missense) | 26093141 | HFE | G | A | 0.0768 | 6.61E-20 | 2.91E-04 | 3.44E-03 | U91328.19 |
| 15 | dMRI Crossing pontine tract | 1 | dMRI_TBSS_MO_Pontine_crossing_tract | 7 | rs2286184 | 84630516 | SEMA3D | C | T | 0.201 | 5.31E-17 | 6.02E-09 | 1.58E-04 |  |
| 16 | dMRI Corpus callosum (genu) | 1 | dMRI_TBSS_OD_Genu_of_corpus_callosum | 7 | rs12113919 | 117612315 | intergenic | C | G | 0.416 | 3.96E-12 | 1.44E-04 | 1.84E-03 | CTTNBP2 |
| 17 | Volume Brain | 2 | volume_MaskVol | 7 | rs2908004 (missense) | 120969769 | WNT16 | G | A | 0.4455 | 3.55E-16 | 7.07E-09 | 2.50E-04 | CPED1, FAM3C |
| 18 | T2* Putamen | 2 | SWI_T2*_putamen_L+R | 8 | rs35469695 | 23406169 | SLC25A37 | C | G | 0.174 | 2.22E-12 | 2.11E-02 | 2.17E-01 | SLC25A37 |
| 19 | Volume Pallidum | 3 | T1_FIRST_pallidum_volume_L+R | 8 | rs2923405 | 42448126 | SMIM19/SLC20A2 | T | G | 0.583 | 3.31E-17 | 1.34E-04 | 5.98E-03 | SMIM19, SLC20A2 |
| 20 | T2* Pallidum | 2 | SWI_T2*_pallidum_L+R | 8 | rs2978098 | 101676675 | SNX31 | A | C | 0.468 | 6.43E-15 | 1.08E-05 | 3.23E-01 | SNX31 |
| 21 | Volume Cerebellum | 3 | T1_FAST_ROIs_L_cerebellum_crus_I | 9 | rs72754248 | 119061396 | PAPPA | G | A | 0.069 | 1.38E-17 | 4.23E-06 | 2.01E-01 |  |
| 22 | T2* Pallidum, putamen and caudate | 17 | SWI_T2*_pallidum_L+R | 10 | rs10764176 (missense) | 18,242,311 | SLC39A12 | A | G | 0.3 | 3.30E-21 | 1.01E-11 | 9.71E-02 | SLC39A12 |
| 23 | T2* Caudate | 3 | SWI_T2*_caudate_L+R | 10 | rs12570727 | 18,425,519 | CACNB2 | G | A | 0.394 | 2.17E-22 | 2.20E-10 | 6.23E-04 | SLC39A12-AS1 |
| 24 | rfMRI Parietal, temporal and prefrontal nodes | 20 | NODEamps100_0002 | 10 | rs2274224 (missense) | 96039597 | PLCE1 | G | C | 0.431 | 6.55E-19 | 1.73E-03 | 7.21E-02 | NOC3L, PLCE1, PLCE1-AS1 |
| 25 | rfMRI Prefrontal nodes | 6 | NODEamps25_0013 | 10 | rs11596664 | 134280157 | INPP5A | C | T | 0.439 | 1.97E-15 | 2.23E-05 | 3.60E-02 | INPP5A RP11, 432J24.6 |
| 26 | T2* Pallidum | 3 | SWI_T2*_pallidum_L+R | 11 | rs11230859 | 61769972 | intergenic | G | A | 0.663 | 2.31E-17 | 6.39E-03 | 4.83E-02 |  |
| 27 | dMRI Crossing pontine tract | 1 | dMRI_TBSS_MO_Pontine_crossing_tract | 11 | rs4935898 (missense) | 124742385 | ROBO3 | G | A | 0.048 | 1.76E-19 | 2.47E-05 | 2.47E-01 |  |
| 28 | Volume Mesencephalon (WM cerebellum, brainstem) | 3 | volume_Right-Cerebellum-White-Matter | 12 | rs4301837 | 102336310 | DRAM1 GNPTAB CHPT1 | T | C | 0.501 | 3.40E-13 | 3.37E-04 | 1.23E-02 | GNPTAB, CHPT1, DRAM1 |
| 29 | Volume Hippocampus | 2 | T1_FAST_ROIs_R_hippocampus | 12 | rs7315280 | 117320938 | intergenic | A | G | 0.115 | 7.06E-14 | 6.80E-05 | 6.69E-01 | FBXW8, HRK |
| 30 | Volume Putamen | 4 | volume_Right-Putamen | 14 | rs945270 | 56200473 | intergenic | C | G | 0.419 | 3.67E-14 | 9.27E-06 | 3.32E-03 |  |
| 31 | Volume and area of precuneus and cuneus | 11 | T1_FAST_ROIs_R_intracalc_cortex | 14 | rs74826997 | 59628609 | DAAM1 | T | C | 0.125 | 2.46E-16 | 3.08E-07 | 2.88E-02 | L3HYPDH, JKAMP |
| 32 | Thickness, area and volume of primary sensorimotor cortex | 15 | a2009s_lh_s_central_area | 15 | rs4924345 | 39639898 | RP11-624L4.1 | A | C | 0.081 | 3.27E-53 | 1.69E-27 | 1.01E-06 |  |
| 33 | Volume 4th ventricle | 1 | volume_4th-Ventricle | 15 | rs2642636 | 58363242 | ALDH1A2 | C | G | 0.415 | 5.24E-16 | 5.63E-03 | 1.81E-01 | ALDH1A2, AQP9 |
| 34 | dMRI Uncinate | 4 | dMRI_ProbtrackX_ISOVF_unc_r | 16 | rs7197215 | 51449978 | intergenic | A | G | 0.566 | 2.24E-15 | 4.50E-02 | 1.43E-04 |  |
| 35 | Volume Cerebellum IX | 2 | T1_FAST_ROIs_L_cerebellum_IX | 17 | rs9905515 | 35261073 | RP11-445F12.1 | G | C | 0.23 | 3.32E-13 | 9.84E-06 | 2.70E-04 |  |
| 36 | T2* Caudate and putamen | 6 | SWI_T2*_putamen_L+R | 17 | rs668799 | 40716235 | COASY | C | T | 0.278 | 1.43E-17 | 1.79E-04 | 9.86E-04 | TUBG2, CNTNAP1, FAM134C, NAGLU, BECN1, HSD17B1, PLEKHH3 |
| 37 | Volume WM lesions | 1 | T2_FLAIR_BIANCA_WMH_volume | 17 | rs3744020 | 73871773 | TRIM47 | G | A | 0.188 | 1.15E-12 | 6.05E-06 | 3.36E-02 | TRIM47, TRIM65, RP11-552F3.9, etc. |
| 38 | dMRI Crossing pontine tract | 1 | dMRI_TBSS_MO_Pontine_crossing_tract | 18 | rs2928990 | 49421125 | intergenic | T | G | 0.898 | 3.97E-16 | 3.96E-05 | 2.27E-03 |  |

**Supplementary Figures 6** and **7** provide genome-wide association plots (also known as Manhattan plots) and QQ-plots for all 3,144 IDPs and the subset of IDPs listed in **Table 1**, respectively. Having identified a SNP as being associated with a given IDP, it can be useful then to explore the association with all other IDPs via a PheWAS (Phenome Wide Association Study) plot. **Supplementary Figure 8** shows the PheWAS plots for all 78 SNPs listed in **Supplementary Table 6** with -log10p>11. The Oxford Brain Imaging Genetics (BIG) web browser (see **URLs**) allows researchers to view the PheWAS for any SNP of interest. We found that 4 of the 78 SNPs were associated (p-value < 0.05/3144, i.e., -log_10_ p-value > 4.79) with all 3 classes of structural, dMRI and functional measures, and these were all SNPs in cluster 31 of **Table 1** (see pages 62-65 of **Supplementary Figure 8**. This genetic locus is associated with the volume of the precuneus and cuneus, dMRI measures for the forceps major (a fibre bundle connecting left and right cuneus), and two functional connections (parcellation 100 edges 614 and 619, which connect the precuneus to other cognitive networks). **Supplementary Figure 9** illustrates the sharing of association signal across IDPs at the 615 unique SNPs listed in **Supplementary Table 5. Supplementary Figure 10** shows the relationship between the number of associations found and the estimated SNP heritability for each IDP.

Overall, our results clearly replicate the majority of the loci identified by the ENIGMA consortium in two separate GWAS of 7 brain subcortical volume IDPs in up to 13,171 subjects^7^, and of hippocampal volume in 33,536 subjects (although not all reached genome-wide significance, likely due to the smaller sample size in our study: **Supplementary Figure 11**). We also replicate an association between volume of white matter hyperintensities (“lesions”) and SNPs in *TRIM47* (e.g., rs3744017, P=1.4E-12, cluster 37) ^18^

It can be challenging to precisely interpret the function of SNPs identified in GWAS. We find that most of the SNPs in the 38 loci in **Table 1** are either in genes, including 7 missense SNPs and 2 SNPs in untranslated regions (UTRs), or in high linkage disequilibrium (LD) with SNPs that are themselves in the genes of interest, and many are significant expression quantitative trait loci (eQTLs) in the GTEx database ^19^ In total we find 17 genetic loci that can be linked to genes that broadly contribute to brain development, patterning and plasticity (out of the 38 clusters reported in **Table 1**; for more details, see **Supplementary Note 3**). In what follows we focus on some of the most compelling examples.

A major source of cross-subject differences seen in T2* data is microscopic variations in magnetic field, often associated with iron deposition in ageing and pathology ^20^ We identified many associations between T2* measurements in the caudate, putamen and pallidum and SNPs in genes (*TF*, rs4428180, P_min_=2.23E-22, cluster 7; *HFE*, rs1800562 (missense) P_min_=6.6E-20, cluster 14; *SLC25A37*, rs35469695, P_min_=2.22E-12, cluster 18) or near genes (*FTH1*, rs11230859, P_min_=2.31E-17, cluster 26) known to affect iron transport and storage, as well as neurodegeneration with brain iron accumulation (NBIA)^21^ (*COASY*, rs668799, P_min_=1.43E-17, cluster 36). In particular, 22 a SNP in *HFE* (s1800562) is associated with haemoglobin levels, iron status 23 24 biomarkers and LDL cholesterol. In addition to *TF*, which transports iron from the intestine, and *SLC25A37*, a mitochondrial iron transporter, we identified four further SNPs that are either coding SNPs for, or eQTLs of, genes involved in transport of nutrients and minerals: *SLC44A5* (rs76934732, P=8.51E-13, cluster 1), *SLC39A8/ZIP8* (rs13107325 (missense) P_min_=1.04E-42, cluster 10), *SLC20A2* (rs2923405, P_min_=3.31E-17, cluster 19) and *SLC39A12/ZIP12* (rs10764176 (missense), P_min_=3.3E-21, cluster 22).

Interrogating images at a voxel-wise level can provide further insight about detailed spatial localisation of SNP associations (e.g., a specific thalamic nucleus), as well as possibly identifying additional associated areas not already well captured by the IDPs (while keeping in mind the statistical dangers of potential circularity^25^). For instance, by looking at the difference between the average T2* image from the subjects having 0 vs. 1 copy of the rs4428180 *(TF)* non-reference allele, effects of this SNP were found not just in the putamen and pallidum, but also in additional, much smaller or more localised regions of subcortical structures that were not included as IDPs (**Figure 2**). We similarly created in **Figure 2** the voxelwise differences associated with 3 additional SNPs, from the most significant GWAS associations with T2* in the putamen as seen in the Manhattan plot. This approach also allowed us to observe grey matter volume effects across the entire brain associated with rs13107325 (*SLC39A8*/*ZIP8*) (**Figure 3**), which has been linked in many previous (non-imaging) GWAS to e.g., intelligence^26^, schizophrenia^27^, blood pressure^28^ and higher risk of cardiovascular death^29^. These effects could now be observed in a very relevant brain region, the anterior cingulate cortex, which is well-known for its multifaceted roles including precisely in fluid intelligence, schizophrenia and in modulating autonomic states of cardiovascular arousal^32^.

**Figure 2:**
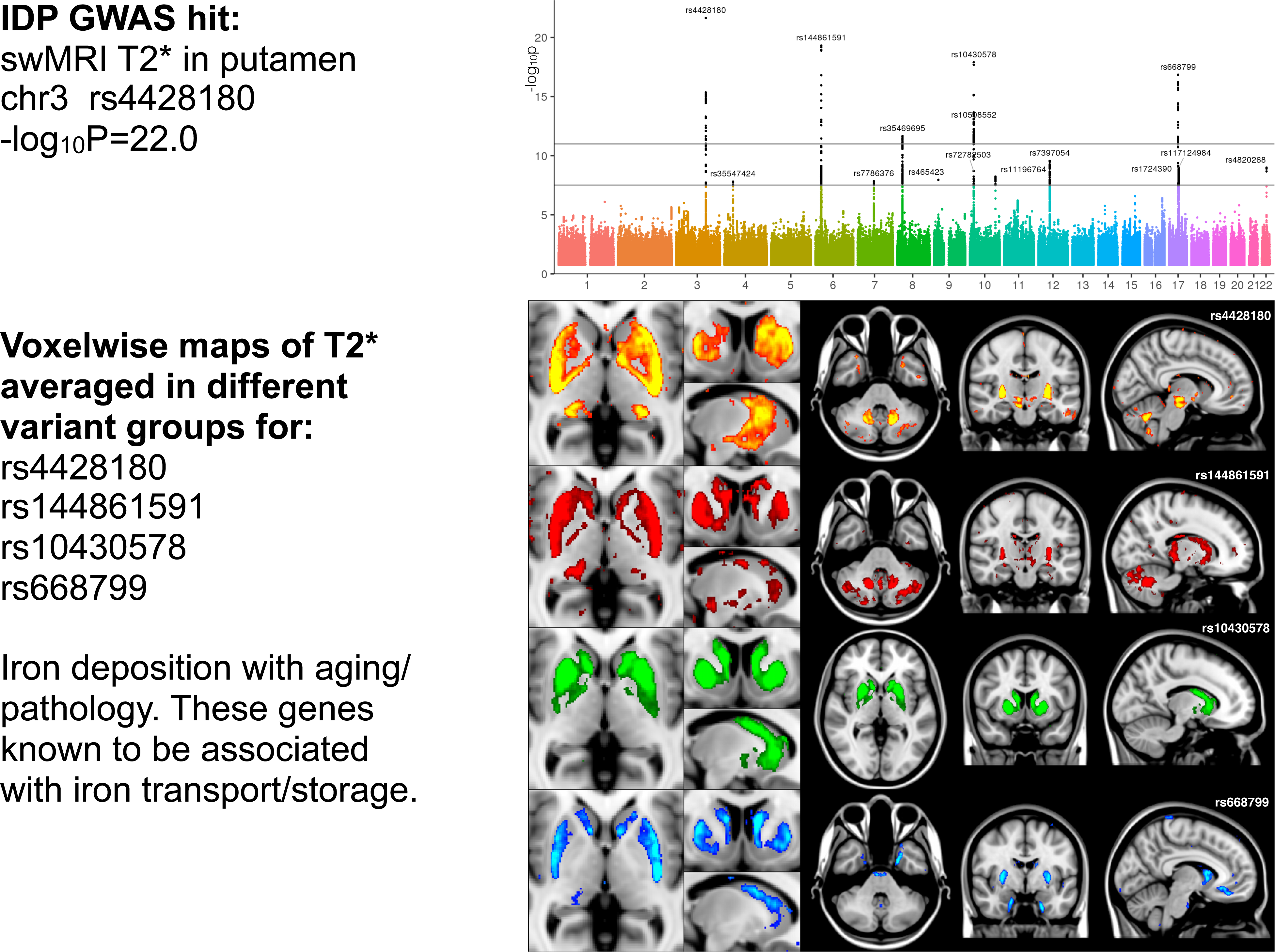
Manhattan plot and spatial mapping of the associations between T2* in the putamen and 4 SNPs. The Manhattan plot relates to the original GWAS for the IDP T2* in the bilateral putamen. The spatial maps show that the 4 SNPs most strongly associated with T2* in the putamen have distinct voxelwise patterns of effect across the whole brain: rs4428180 *(TF)* effect is found in the dorsal putamen and body of the caudate nucleus, but also in the right subthalamic nucleus and substantia nigra, the red nucleus, lateral geniculate nucleus of the thalamus and the dentate nucleus; rs144861591 (*HFE*) in the dorsal striatum, subthalamic nucleus, dentate nucleus and Crus I/II of the cerebellum; rs10430578 *(ZIP12)* in the whole dorsal striatum and pallidum; and rs668799 (*COASY*) in the whole dorsal striatum, subgenual cingulate cortex and entorhinal cortex. The standard MNI152 T1 image is used as background for the spatial maps (left is right). All group difference images (color overlays) are thresholded at a T2* difference of 0.6ms.

**Figure 3:**
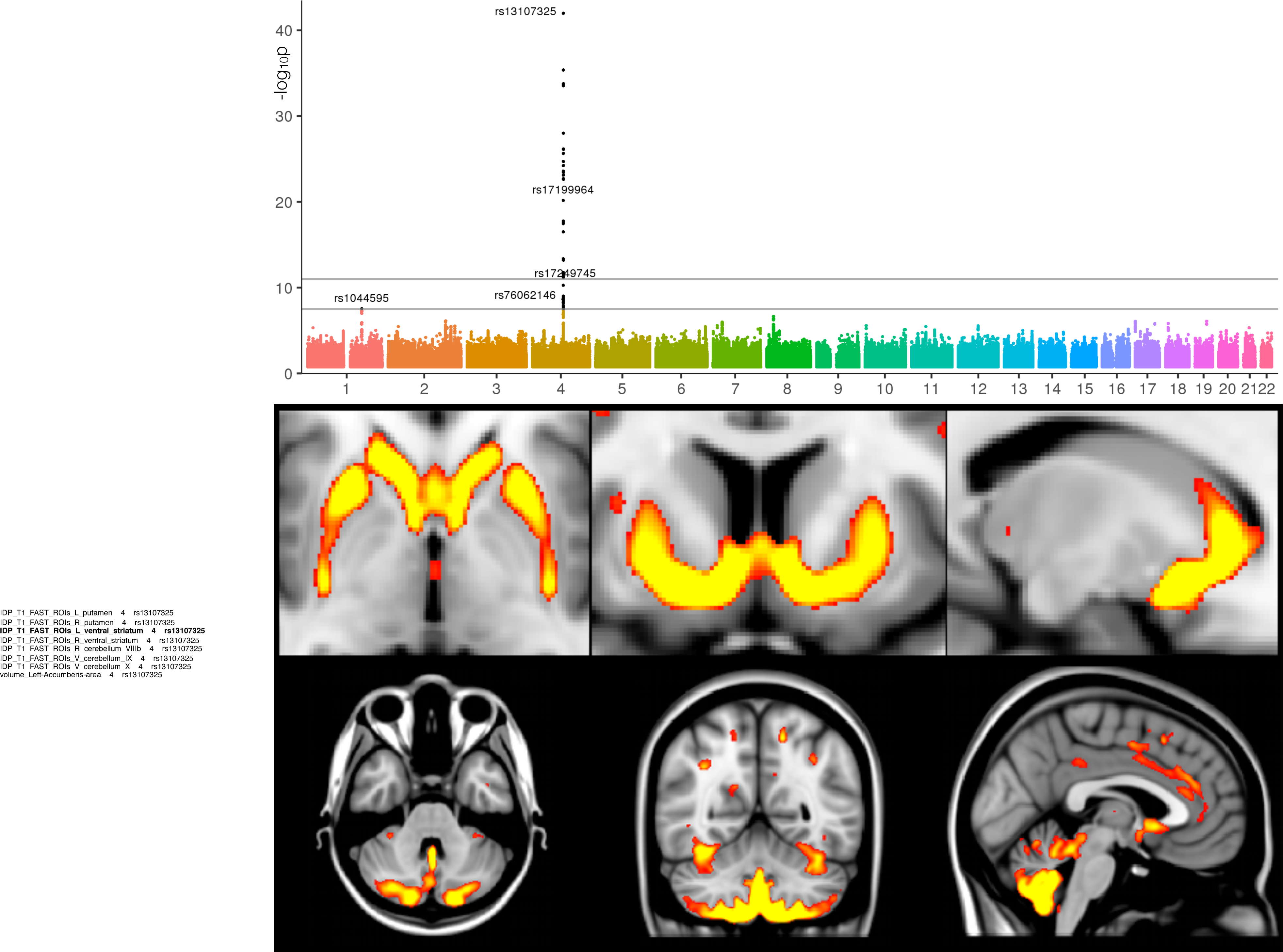
Manhattan plot and spatial mapping of the associations between GM volume and rs13107325 *(SLC39A8/ZIP8).* The Manhattan plot relates to the original GWAS for the IDP of GM volume in the left ventral striatum. The images show spatial mapping of rs13107325 against voxelwise local grey matter volume (GM was averaged across all 1,181 subjects with 1 copy of the non-reference allele, and the average from all 7,215 subjects having 0 copies was subtracted from that, for display in color here; the difference was thresholded at 0.015 - unitless relative measure of local grey matter volume). The maps show that the rs13107325 (SLC39A8/ZIP8) effect is found more generally bilaterally in the ventral caudate, putamen, ventral striatum, anterior cingulate cortex, and with a strong cerebellar contribution (lobules VI-X), particularly in the prefrontal-projecting Crus I/II, which are selectively expanded in humans.

Interestingly, three SNPs related to our white matter IDPs were in genes or eQTLs of genes coding for three proteins of the extracellular matrix (ECM). The first SNP (rs2365715, P=5.38E-12, cluster 2), an eQTL of *BCAN*, is associated with one dMRI microstructural measure in the genu of the corpus callosum. The second SNP (rs3762515, P=4.27E-13, cluster 3), in the 5’ UTR of *EFEMP1*, is associated with the volume of white matter lesions. Finally, the third SNP (rs67827860, P_min_=4.06E-37, cluster 11, **Figure 4**), located in an intron of *VCAN*, is in a cluster associated with multiple dMRI measures of most of the brain white matter tracts (199 IDPs in total). *BCAN* and *VCAN* in particular both code for chondroitin sulfate proteoglycans of the ECM, which are especially crucial for synaptic plasticity^33^ and myelin repair^34^. *VCAN* is, for instance, increased in association with astrocytosis in multiple sclerosis^35^, while both *BCAN* and *VCAN* are differentially regulated following spinal cord injury^36^. *BCAN, EFEMP1* and *VCAN* have been further associated in three separate GWAS with stroke^37^, site of onset of amyotrophic lateral sclerosis^38^ and major 39 depressive disorder, respectively. Furthermore, *EFEMP1* is characterised by tandem arrays of epidermal growth factor (EGF)-like domains, and we also identified a strong association between the whole of the corpus callosum (genu, body and splenium) and a SNP in *HBEGF* (rs4150221, P_min_=8.43E-20, cluster 13), a heparin-binding EGF-like growth factor. Similarly to *BCAN* and *VCAN, HBEGF* plays an important role in oligodendrocyte development and helps recovering WM injury in preterm babies^40^. Remarkably, this means that the vast majority of forebrain WM-related dMRI IDPs are associated in this study with SNPs related to genes coding for proteins involved either in the extracellular matrix, the epidermal growth factor, or both.

**Figure 4:**
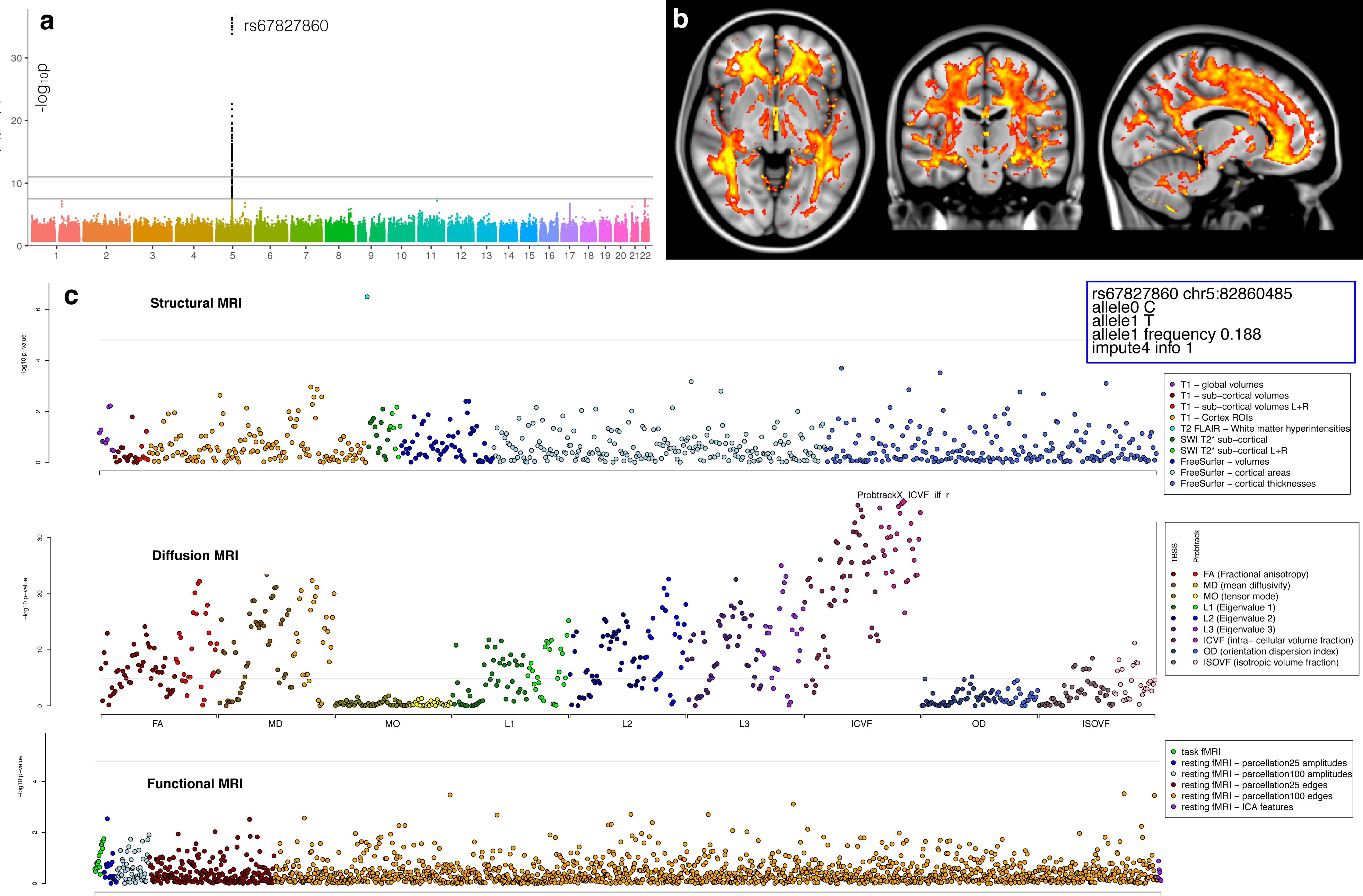
Manhattan plot, spatial mapping and PheWAS plot relating to the association between the dMRI intra-cellular volume fraction (ICVF) measure and rs67827860 *(VCAN).* **a)** The Manhattan plot relates to the original IDP GWAS with the strongest association (ICVF in the right inferior longitudinal fasciculus using tractography, associated with rs67827860). The ICVF parameter, estimated from the NODDI modelling^88^, aims to quantify predominantly intra-axonal water in white matter, by estimating where water diffusion is restricted. **b**) Spatial mapping of rs67827860 against voxelwise ICVF in white matter (ICVF was averaged across all 4,957 subjects with 0 copies of the non-reference allele, and the average from all 2,304 subjects having 1 copy was subtracted from that, for display in color here; the difference was thresholded at 0.005 - unitless fractional measure). Unlike the previous examples of (spatially) very focal effects in T2* and grey matter volume in **Figure. 2 and 3**, the effects of this SNP are extremely widespread across most of the white matter tracts (associated with 45 out of the 199 IDPs in cluster 11, **Supplementary Table 5**)**. c**) The PheWAS plot for SNP rs67827860 shows the association (-log10 p-value) on the y-axis for the SNP rs67827860 with each of the 3,144 IDPs. The IDPs are arranged on the x-axis in the three panels: (top) Structural MRI IDPs, (middle) Structural connectivity dMRI IDPs, (bottom) functional MRI IDPs. Points are coloured to delineate subgroups of IDPs and detailed in the legends. Summary details of SNP rs67827860 are given in the top right box. The grey line shows the Bonferroni multiple testing threshold of 4.79. In addition to the IDP of WM hyperintensities volume, there is a notable association with numerous dMRI IDPs (especially diffusion tensor-derived measures of FA, MO and 1^st^/2^nd^/3^rd^ eigenvalues of the diffusion tensor, as well as additional ICVF measures).

Two additional examples further illustrate highly meaningful correspondences between locations of our brain IDPs and significantly associated genes. First, the volume of the 4^th^ ventricle, which develops from the central cavity of the neural tube, was found to be significantly associated with a SNP in, and eQTL of, *ALDH1A2* (rs2642636, P=5.2E-16, cluster 33). This gene codes for an enzyme which facilitates posterior organ development and prevents human neural tube defects, including spina bifida^41^. Second, we found two SNPs associated with dMRI IDPs of the crossing pontine tract (the part of the pontocerebellar fibre bundle arising from pontine nuclei that decussate across the brain midline to project to contralateral cerebellar cortex) in genes that regulate axon guidance and fasciculation during development (*SEMA3D*, rs2286184, P=5.31E-17, cluster 15 and *ROBO3*, rs4935898 (missense), P=1.76E-19, cluster 27, **Figure 5**). The exact location of our IDP in the crossing fibres of the pons remarkably coincides with the function of *ROBO3*, which is specifically required for axons to cross the midline in the hindbrain (pons, medulla oblongata and cerebellum); mutations in *ROBO3* result in horizontal gaze palsy, a disorder in which the corticospinal and somatosensory axons fail to cross the midline in the medulla^42^. Notably, all three significant associations with the IDP of the crossing pontine tract were found using the mode of anisotropy (MO), which is a tensor-model dMRI measure particularly sensitive to regions of crossing fibres^43^.

**Figure 5:**
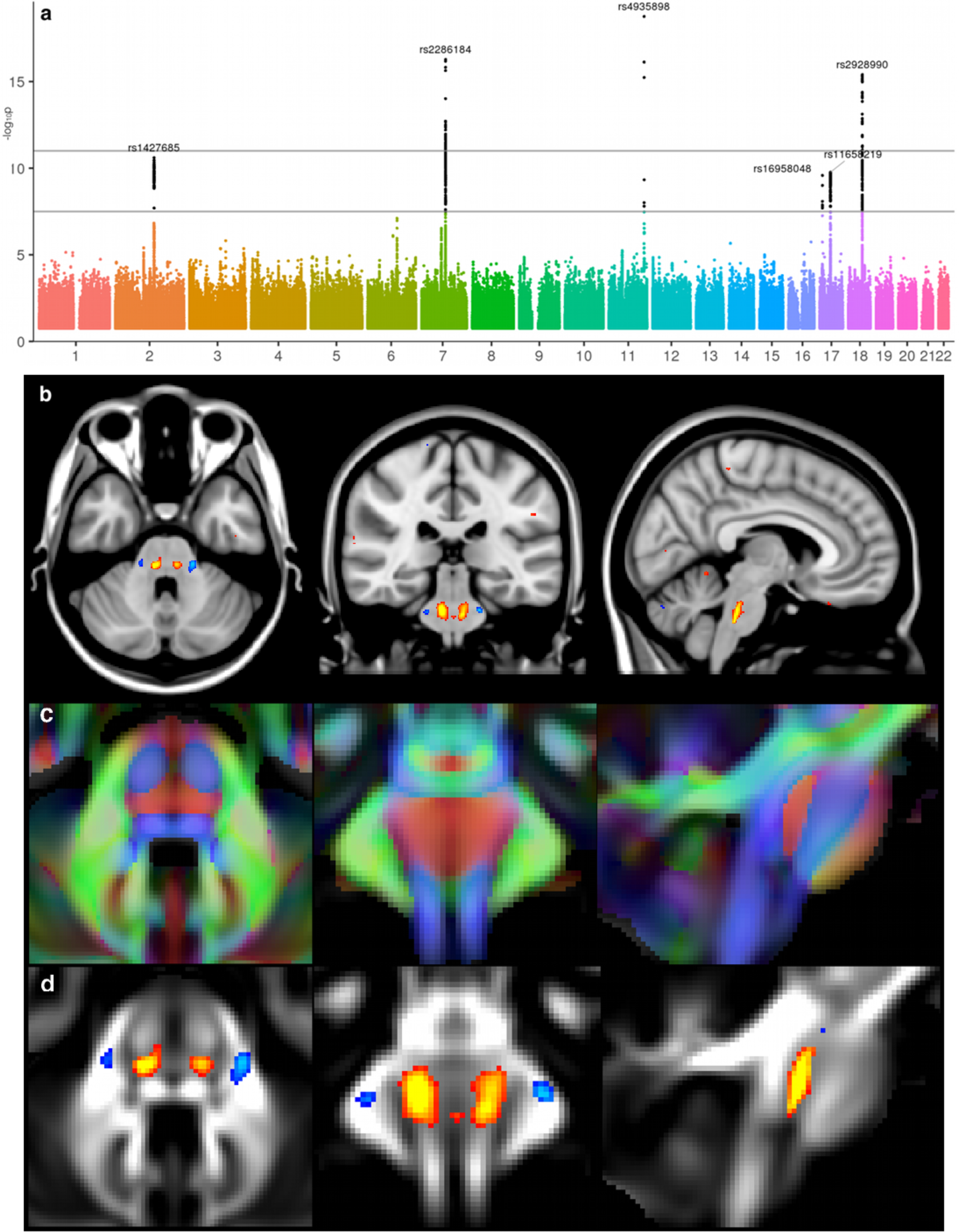
Manhattan plot and spatial mapping of the association between the dMRI tensor mode (MO) measure and SNP rs4935898 *(ROBO3).* The Manhattan plot relates to the original GWAS for the IDP of MO in the crossing pontine tract associated with rs4935898. MO was averaged across all 6,807 subjects with ~0 copy of the non-reference allele, and the average from all 703 subjects having ~1 copy was subtracted from that, for display in red-yellow/blue-lightblue here, thresholded at 0.05 (**b,d**). In (**b**) results are shown overlaid on the MNI152 T1 structural image; in contrast, background image in (**c, d**) is the UK Biobank average FA (fractional anisotropy) that shows clear tract structure within the brainstem. In (**c**) is superimposed the orientat 1196 ion of the fibre tracts (in red, running along the x-axis). The1197 spatial distribution (not shown) for the effects of rs2286184 (*SEMA3D*) on MO is1198 almost identical to that of rs4935898, being again extremely spatially specific, with1199 no extended effect elsewhere in the brain.

### Multi-phenotype association tests

One alternative strategy for analysing large numbers of IDPs is to use multi-trait tests that fit joint models of associations to groups of IDPs. Such approaches can utilise estimates of genetic correlation to boost power. In addition, by analysing *P* traits in one GWAS, these tests can avoid the need to correct for multiple genome-wide scans. We used a multi-trait test (see **Methods**) to analyse 23 groups of IDPs with up to 243 IDPs per group. These IDP groupings were chosen to cover the majority of the IDP classes with significant IDP correlations in each grouping (**Supplementary Table 7**). **Supplementary Figure 12** shows the Manhattan plots for these genome-wide scans. Overall across these 23 groups, we found 278 SNPs at ~160 loci associated with - log_10_ p-value > 7.5 (see **Supplementary Table 8**). We found that 170 of these 278 SNPs survived a correction for 23 scans with -log10 p-value > 8.86 and 138 of these 170 SNPs had a p-value < 0.05 in the larger replication set of 3,456 samples. There can be quite large differences in p-values between the multi-trait tests and the individual IDP tests (**Supplementary Figure 13**), especially when taking account of the smaller number of tests carried out by the multi-trait approach (**Supplementary Figure 14**). We found 25 loci that showed both a significant and replicated multi-trait association for an IDP group, while showing no genome-wide significance in the flanking region for any individual IDP in the corresponding grouping (**Supplementary Table 9**).

Three of these loci show associations with the dMRI MO measures (rs62073157, P=4.07E-11; rs35884657, p=1.04E-09; rs9939914,p=1.15E-11) and all are eQTLs of microtubule related genes *MAPT, TUBA1B* and *TUBB3* respectively. The first SNP rs62073157 resides in a long stretch of LD (43.4-44.9Mb) on chromosome 17 known to contain a common inversion polymorphism^44^. This extended *MAPT* (encoding for Microtubule Associated Protein Tau) region has been repeatedly associated with several neurodegenerative disorders, such as Alzheimer’s disease, where it has been shown to modulate the age of onset ^45^ and to be associated with *APOE* ε4-alleles^46^, fronto-temporal dementia^47^ and progressive supranuclear palsy^48^. Notably, a locus in this *MAPT* region also shows overlap between Alzheimer’s and Parkinson’s disease^49^. Mutations in tubulin genes have been shown to correlate strongly with multiple cortical and subcortical abnormalities^50^.

Another example of the value of the multi-trait testing can be seen in the association between IDPs of global brain volume measurements and a SNP located between *BANK1* and *ZIP8*, previously identified in a GWAS of schizophrenia^51^ (rs35518360, P=4.07E-12). This locus is also part of a multi-modal cluster from our single-trait GWAS that includes subcortical and cerebellar grey matter volumes, pallidum T2* and dMRI in midbrain white matter tracts (cluster 10 in **Supplementary Table 6**). The multi-trait test thus made it possible to uncover this additional association between global brain volume measurement and this locus, which might prove relevant in better understanding observations of smaller brain volume in (first episode/drug-naïve) schizophrenic patients ^52^

Another locus (rs112651271, p=3.23E-11) is associated with a dMRI IDP group encompassing all measurements collected in major white matter tracts. This SNP lies 150Kb upstream of *EDNRA*, which plays a role in potent and long-lasting vasoconstriction, and (likely related to this), has been linked to hypertension and migraine, as well as intracranial aneurysm ^53^.

The multi-trait analysis also uncovered an association with SNPs in the *IL34* gene (rs12928124, p=1.31E-10) and Freesurfer brain volume IDPs. IL-34 is a ligand of the CSF-1 receptor (CSF-1R) that regulates CNS microglial development and has been shown to regulate cortical development in mice ^54^. Il-34 has also been shown to promote clearance of soluble oligomeric amyloid-β, which mediates synaptic dysfunction and neuronal damage in Alzheimer’s disease. ^55^.

### Iron, cardiovascular traits and brain development in brain disorders

Of those genes involved in neurodegenerative disorders which we identified in our single-IDP association analysis, interestingly most mainly code for iron-related proteins. While *TF* and *HFE* might play a relevant role for iron mobilisation and regulation in neurodegenerative disorders such as Parkinson’s disease, Creutzfeldt-Jakob disease, amyotrophic lateral sclerosis and Alzheimer’s disease^56,57^, *SLC25A37* shows increased expression in Alzheimer’s and Friedreich’s ataxia^58^ and mutations in *COASY* are associated with neurodegeneration with brain iron accumulation ^21^.

One notable exception, is in an LD region encompassing significant SNPs in both *MRC1* and *ZIP12* (cluster 22), which has been linked to neurodegenerative/neuropsychiatric disorders and cardiovascular processes (as opposed to iron-related processes). SNPs in *MRC1* have been shown in a GWAS to be associated with risk of cardiovascular disease ^59^ and *MRC1* expression is increased in a model of Alzheimer’s disease ^60^, while *ZIP12* demonstrates altered expression in the cortex of subjects with schizophrenia ^61^. Our significant SNPs in *ZIP8* (cluster 10) show a similar overlap, and *ZIP8* hit has been found associated both with schizophrenia and Parkinson’s disease^62^, as well cardiovascular death^29^.

Similarly to *ZIP8* and *ZIP12*, of those genes related to mental health disorders identified both in the single-IDP and multi-trait analyses, most are strongly involved in brain development and plasticity. This is the case of *VCAN*, for which SNPs have been associated in a GWAS with major depressive disorder ^39^, *SEMA3D* and *DAAM1*, which might both contribute to schizophrenia ^63,64^, *ROBO3* that may be associated with autism^65^ and *CTTNBP2*, for which disruption is related to autism^66^, and knockdown reduces the density and size of dendritic spines in neurons (rs12113919, eQTL of *CTTNBP2*, P=3.96E-12, cluster 16). This latter SNP was interestingly associated here with one dMRI measures in the corpus callosum, a white matter tract that has been shown in dMRI meta-analyses to be the most consistently disrupted white matter tract in autism^67,68^.

### Genetic correlation with neurodegenerative, psychiatric and personality traits

We measured the genetic correlation (hence also co-heritability) between a subset of heritable IDPs and 10 neurodegenerative, psychiatric and personality traits (see **Methods**). We found some suggestive evidence of elevated levels of non-zero genetic correlation for amyotrophic lateral sclerosis (ALS), schizophrenia and stroke, mainly with dMRI measures in white matter tracts (**Supplementary Figure 15**). The strongest genetic correlation for ALS (P<10^−3^) was found in the genu of the corpus callosum (with a co-heritability of 0.63). This result is in line with consistent findings of corpus callosum involvement in this degenerative disorder ^69^. Correlations found in schizophrenia with the tapetum (P<10^−3^) were likely due to partial volume effects, given that the next most strongly associated IDPs reflect ventricular and thalamic volume, which are some of the most robust volumetric findings in this mental health disorder ^52^; hence it is interesting to see the genetic input into this volumetric disease association. While more modest correlations in stroke were observed, it was across a wide range of dMRI IDPs, with the strongest genetic correlations (P<10^−2^) in the corona radiata, internal capsule and thalamic radiations, i.e., white matter tracts that follow the probabilistic distribution of stroke ^70^. **Supplementary Table 10** contains genetic correlation estimates for all IDP/trait combinations with nominal p-value < 0.01, to highlight which IDPs occur in the tails of these distributions. However, in line with previous observations [Bulik-Sullivan 2015], we also found evidence that the LDSCORE regression approach^71^ for estimating genetic correlation seems best suited to pairs of traits both of which are heritable and polygenic in genetic aetiology. For example, the deflated p-value distribution for the correlation of IDPs with Alzheimer’s is driven by the large *APOE* association for Alzheimer’s disease on chromosome 19.

### Partitioning heritability by functional annotation

We applied a statistical approach that partitions the additive genetic heritability of a set of common variants for each of the 3,144 IDPs according to 24 functional annotations of the genome^71^. **Figure 6** summarizes which functional annotations show enrichment stratified by 23 groups of IDPs (see also **Supplementary Table 11**). We find that regions of the genome annotated as Super Enhancers and several histone modifications show enrichment across many of the structural and diffusion IDP groups. Regions of the genome enriched for histone modification H3K27me3 (and indicating strong evidence for silenced genes) show depletion of heritability across many of the IDP classes (**Supplementary Figure 16**). IDP groups such as T1 subcortical volumes, dMRI FA and ICVF show the strongest evidence of enrichment across multiple categories. The resting fMRI connectivity edge IDPs show no elevated enrichment, consistent with these traits showing low overall levels of heritability (**Figure 1**). **Supplementary Figure 17** provides the results of this partitioning analysis for each IDP.

**Figure 6:**
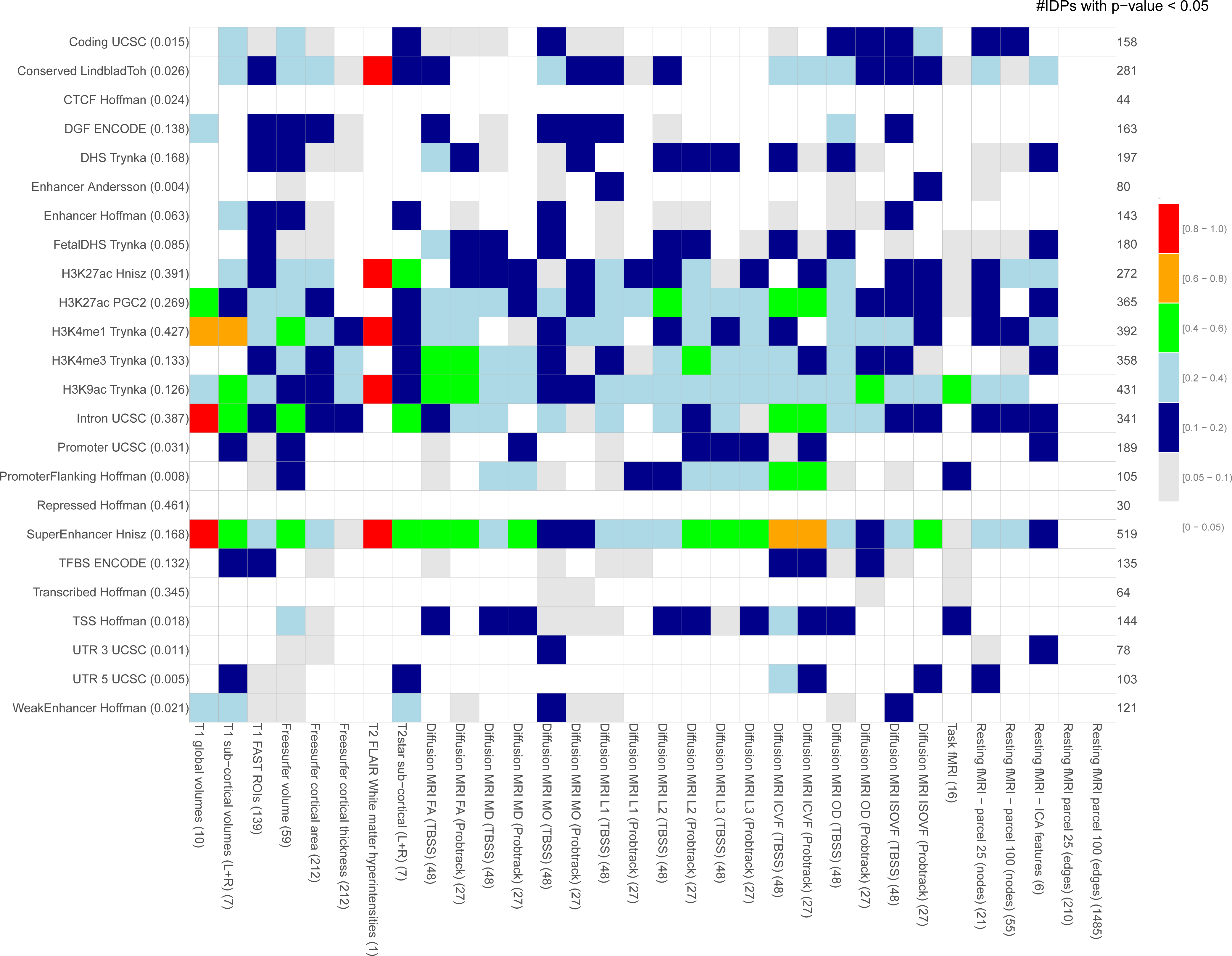
Partitioning of heritability by functional category. The plot shows the 1202 proportion of IDPs in each of the 23 IDP groupings (x-axis) that show a nominal 1203 *enrichment* p-value < 0.05 for the 24 functional categories (y-axis). The total number 1204 of such IDPs for each category is given on the right edge of the plot. The number of 1205 IDPs in each IDP group is listed in brackets in the x-axis labels. The proportion of the 1206 genome annotated by each functional category is listed in brackets in the y-axis labels.

## Conclusions

Bringing together researchers with backgrounds in brain imaging and genetic association was key to this work. We have uncovered a large number of associations at the nominal level of GWAS significance (-log10 p-value > 7.5) and at a more stringent threshold (-log10 p-value > 11) designed to (probably over-conservatively) control for the number of IDPs tested. Our use of multi-trait tests uncovered further novel loci. We find associations with all the main IDP groups except the task fMRI measures (despite these measures containing usable signal, for example having unique and strong cognitive associations^4^). We mainly found associations between our MRI measures and genes involved in brain development and plasticity, as well as with genes contributing to transport of nutrients and minerals. Most of these genes have also been demonstrated to contribute to a vast array of disorders including major depression disorder, cardiovascular disease, schizophrenia, amyotrophic lateral sclerosis and Alzheimer’s disease. We further uncovered enrichments of functional annotations for many of the structural and diffusion IDPs.

A valuable aspect of this work has been to link the associated SNPs back to spatial properties of the voxel-level brain imaging data. For example, we have linked SNPs associated with IDPs to both highly spatially localized (**Figure. 2,3,5**) and widely spatially distributed (**Figure 4**) effects, restricting these voxelwise analyses to the same imaging modality from which the original phenotypic association was found (though of course other modalities could also be tested in the same way). In addition, looking at PheWAS plots has been useful when working with so many phenotypes. It has allowed investigation of the overall patterns of association and has led to the identification of SNP associations that span multiple modalities.

We have used two separate sets of 930 and 3,456 samples to replicate a large number of the associations uncovered at the discovery phase. Over the next few years, the number of UK Biobank participants with imaging data will gradually increase to 100,000, which will allow a much more complete discovery of the genetic basis of human brain structure, function and connectivity. Combining the discovery and replication samples will likely also lead to novel associations, as will the use of methods that can analyze the huge IDP × SNP matrix of summary statistics of association. A potential avenue of research will involve attempting to uncover causal pathways that link genetic variants to IDPs and then onto a range of neurological, psychiatric and developmental disorders.

## Methods

### Imaging data and derived phenotypes

The UK Biobank Brain imaging protocol consists of 6 distinct modalities covering structural, diffusion and functional imaging, summarised in **Supplementary Table 1**. For this study, we primarily used data from the February 2017 release of ~10,000 participants’ imaging data (and an additional ~5,000 subjects’ data released in January 2018 provided the larger replication sample).

The raw data from these 6 modalities has been processed for UK Biobank to create a set of imaging derived phenotypes (IDPs)^4,72^. These are available from UK Biobank, and it is these IDPs from the 2017/18 data releases that we used in this study.

In addition to the IDPs directly available from UK Biobank, we created two extra sets of IDPs. Firstly, we used the FreeSurfer v6.0.0 software^73,74^ to model the cortical surface (inner and outer 2D surfaces of cortical grey matter), as well as modelling several subcortical structures. We used both the T1 and T2-FLAIR images as inputs to the FreeSurfer modelling. FreeSurfer estimates a large number of structural phenotypes, including volumes of subcortical structures, surface area of parcels identified on the cortical surface, and grey matter cortical thickness within these areas. The areas are defined by mapping an atlas containing a canonical cortical parcellation onto an individual subject’s cortical surface model, thus achieving a parcellation of that surface. Here we used two atlases in common use with FreeSurfer: the Desikan-Killiany-Tourville atlas (denoted “DKT” ^75^) and the Destrieux atlas (denoted “a2009s” ^76^). The DKT parcellation is gyral-based, while Destrieux aims to model both gyri and sulci based on the curvature of the surface. Cortical thickness is averaged across each parcel from each atlas, and the cortical area of each parcel is estimated, to create two IDPs for each parcel. Finally, subcortical volumes are estimated, to create a set of volumetric IDPs.

Secondly, we applied a dimension reduction approach to the large number of functional connectivity IDPs. Functional connectivity IDPs represent the network “edges” between many distinct pairs of brain regions, comprising in total 1,695 distinct region-pair brain connections (see **URLs**). In addition to this being a very large number of IDPs from which to interpret association results, these individual IDPs tend to be significantly noisier than most of the other, more structural, IDPs. Hence, while we did carry out GWAS for each of these 1,695 connectivity IDPs, we also reduced the full set of connectivity IDPs into just 6 new summary IDPs using data-driven feature identification. We did this dimensionality reduction by applying independent component analysis (ICA^77^), applied to all functional connectivity IDPs from all subjects, to find linear combinations of IDPs that are independent between the different features (ICA components) identified^78^. We carried out the ICA feature estimation without any use of the genetic data, and we maximized independence between component IDP weights (as opposed to subject weights). We used split-half reproducibility (across subjects) to optimize both the initial dimensionality reduction (14 eigenvectors from a singular value decomposition was found to be optimal) and also the final number of ICA components (6 ICA components was optimal, with reproducibility of ICA weight vectors greater than r=0.9). The resulting 6 ICA features were then treated as new IDPs, representing 6 independent sets (or, more accurately, linear combinations) of the original functional connectivity IDPs. These 6 new IDPs were added into the GWAS analyses. The 6 ICA features explain 4.9% of the total variance in the full set of network connection features, and are visualized in **Supplementary Figure 18**. More details of the ICA analysis of the resting state data, together with browsing functionality of the highlighted brain regions can be found on the FMRIB Biobank Resource web page (see **URLs**).

We organised all 3,144 IDPs into 9 groups (**Supplementary Table 12**), each having a distinct pattern of missing values (not all subjects have usable, high quality data from all modalities^4^). For the GWAS in this study we did not try to impute missing IDPs due to low levels of correlation observed across groups.

The distributions of IDP values varied considerably between phenotype classes, with some phenotypes exhibiting significant skew (**Supplementary Figure 19**) which would likely invalidate the assumptions of the linear regression used to test for association. To ameliorate this we quantile normalized each of the IDPs before association testing. This transformation also helps avoid undue influence of outlier values. We also (separately) tested an alternative process in which an outlier removal process was applied to the un-transformed IDPs; this gave very similar results for almost all association tests, but was found to reduce the significance of a very small number of associations. This possible alternative method for IDP “preprocessing” was therefore not followed through (data not shown).

### Genetic data processing

We used the imputed genetic dataset made available by UK Biobank in its July 2017 release^6^. This consists of >92 million autosomal variants imputed from the Haplotype Reference Consortium (HRC) reference panel^79^ and a merged UK10K + 1000 Genomes reference panel. We first identified a set of 12,623 participants who had also been imaged by UK Biobank. We then applied filters to remove variants with minor allele frequency (MAF) below 0.1% and with an imputation information score below 0.3, which reduced the number of SNPs to 18,174,817. We then kept only those samples (subjects) estimated to have recent British ancestry using the sample quality control information provided centrally by UK Biobank^6^ (using the variable *in.white.British.ancestry.subset* in the file *ukb_sqc_v2.txt);* population structure can be a serious confound to genetic association studies^80^, and this type of sample filtering is standard. This reduced the number of samples to 8,522. The UK Biobank dataset contains a number of close relatives (3^rd^ cousin or closer). We therefore created a subset of 8,428 nominally unrelated subjects following similar procedures in Bycroft et al. (2017). After running GWAS on all the (SNP) variants in the 8,428 samples we applied three further variant filters to remove variants with a HWE (Hardy-Weinberg equilibrium) p-value less than 10^−7^, remove variants with MAF<0.1% and to keep only those variants in the HRC reference panel. This resulted in a dataset with 11,734,353 SNPs.

We used two separate datasets for replicating the associated variants found in this study. The first set of 930 samples were a subset of the 1,279 samples with imaging data that we did not use for the main GWAS, which had been primarily excluded due to not being in the recent British ancestry subset. An examination of these samples according the genetic principal components (PCs) revealed that many of those samples are mostly of European ancestry (**Supplementary Figure 20**). We selected 930 samples with a 1^st^ genetic PC < 14 from **Supplementary Figure 20** and these constituted the replication sample. In January 2018 a further tranche of 4,588 samples with imaging data was released by UK Biobank. Of these subjects, we selected 3,956 subjects that both had genetic data available and also were imaged in the same imaging center as the discovery sample. We applied the same pre-processing pipeline as for the discovery set. We then restricted this to 3,456 subjects that were of recent British ancestry and replication tests were then conducted on these 3,456 subjects.

### Potential Confounds for brain IDP GWAS

There are a number of potential confounding variables when carrying out GWAS of brain IDPs. We used three sets of covariates in our analyses relating to (a) imaging confounds (b) measures of genetic ancestry, and (c) non-brain imaging body measures.

We identified a set of variables likely to represent imaging confounds, for example those being associated with biases in noise or signal level, corruption of data by head motion or overall head size changes. For many of these we generated various versions (for example, using quantile normalization and also outlier removal, to generate two versions of a given variable, as well as including the squares of these to help model nonlinear effects of the potential confounds). This was done in order to generate a rich set of covariates and hence reduce as much as possible potential confounding effects on analyses such as the GWAS, which are particularly of concern when the subject numbers are so high.^4,81^

Age and sex are can be variables of biological interest, but can also be sources of imaging confounds, and here were included in the confound regressors. Head motion is summarized from the rfMRI and tfMRI as the mean displacement (in mm) between one timepoint and the next, averaged over all timepoints and across the brain. Head motion can be a confounding factor for all modalities and not just those comprising timeseries of volumes, but is only readily estimable from the timeseries modalities. Nevertheless, the amount of head motion is expected to be reasonably similar across all modalities (e.g., correlation between head motion in resting and task fMRI is *r*=0.52) and so it is worth using fMRI-derived head motion estimates as confound regressors for all modalities.

The exact location of the head and the radio-frequency receive coil in the scanner can affect data quality and IDPs. To help account for variations in position in different scanned participants, several variables have been generated that describe aspects of the positioning (see **URLs**). The intention is that these can be useful as “confound variables”, for example these might be regressed out of brain IDPs before carrying out correlations between IDPs and non-imaging variables. TablePosition is the Z-position of the coil (and the scanner table that the coil sits on) within the scanner (the Z axis points down the centre of the magnet). BrainCoGZ is somewhat similar, being the Z-position of the centre of the brain within the scanner (derived from the brain mask estimated from the T1-weighted structural image). BrainCoGX is the X-position (left-right) of the centre of the brain mask within the scanner. BrainBackY is the Y-position (front-back relative to the head) of the back of brain mask within the scanner.

UK Biobank brain imaging aims to maintain as fixed an acquisition protocol as possible during the 5-6 years that the scanning of 100,000 participants will take. There have been a number of minor software upgrades (the imaging study seeks to minimise any major hardware or software changes). Detailed descriptions of every protocol change, along with thorough investigations of the effects of these on the resulting data, will be the subject of a future paper. Here, we attempted to model any long-term (over scan date) changes or drifts in the imaging protocol or software or hardware performance, by generating a number of data-driven confounds. The first step was to form a temporary working version of the full subjects x IDPs matrix with outliers limited (see below) and no missing data, using a variant of low-rank matrix imputation with soft thresholding on the eigenvalues^82^. Next, the data is temporally regularized (approximate scale factor of several months with respect to scan date) with spline-based smoothing. We then applied PCA and kept the top 10 components kept, to generate a basis set reflecting the primary modes of slowly-changing drifts in the data.

To describe the full set of imaging confounds we use a notation where subscripts “i” indicate quantile normalization of variables, and “m” to indicate median-based outlier removal (discarding values greater than 5 times the median-absolute-deviation from the overall median). If no subscript is included, no normalization or outlier removal was carried out. Certain combinations of normalization and powers were not included, either because of very high redundancy with existing combinations, or because a particular combination was not well-behaved. The full set of variables used to create the confounds matrix are:

- a = age at time of scanning, demeaned (cross-subject mean subtracted)
- s = sex, demeaned
- q = 4 confounds relating to the position of the radio-frequency coil and the head in the scanner (see above), all demeaned
- d = 10 drift confounds (see above)
- m = 2 measures of head motion (one from rfMRI, one from tfMRI)
- h = volumetric scaling factor needed to normalise for head size ^83^

The full matrix of imaging confounds is then:

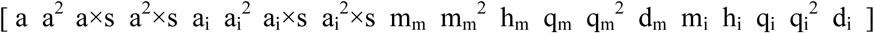

Any missing values in this matrix are set to zero after all columns have had their mean subtracted. This results in a full-rank matrix of 53 columns (ratio of maximum to minimum eigenvalues = 42.6). For additional discussion on the dangers and interpretation of imaging confounds in big imaging data studies, particularly in the context of disease studies, see ^81^.

Genetic ancestry is a well-known potential confound in GWAS. We ameliorated this by filtering out samples that were not of recent British ancestry. However, a set of 40 genetic principal components (PCs) has been provided by UK Biobank^6^ and we used these PCs as covariates in all of our analysis. The matrix of imaging confounds, together with a matrix of 40 genetic principal components, was regressed out of each IDP before the analyses reported here.

There exist a number of substantial correlations between IDPs and non-genetic variables collected on the UK Biobank subjects^4^. Based on this, we also carried out some analyses involving variables relating to Blood Pressure (Diastolic and Systolic), Height, Weight, Head Bone Mineral Density, Head Bone Mineral Content and 2 principal components from the broader set of bone mineral variables available (see **URLs**). **Supplementary Figure 21** shows the association of these 8 variables against the IDPs and shows significant associations. These are variables that likely have a genetic basis, at least in part. Genetic variants associated with these variables might then produce false positive associations for IDPs. To investigate this, we ran GWAS for these 8 traits (conditioned on the imaging confounds and genetic PCs) (**Supplementary Figure. 22**). We also ran a parallel set of IDP GWAS with these “body confounds” regressed out of the IDPs.

### Heritability and genetic correlation of IDPs

We used a linear mixed model implemented in the SBAT (Sparse Bayesian Association Test) software (see **URLs**) to calculate additive genetic heritabilities for the *P*=3,144 traits. To estimate genetic correlations we used a multi-trait mixed model. If *Y* is an *N×P* matrix of *P* phenotypes (columns) measured on *N* individuals (rows) then we use the model

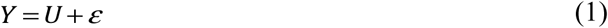

where *U* is an *N×P* matrix of random effects and *ε* is a *N×P* matrix of residuals and these are modelled using Matrix normal distributions as follows

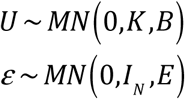

In this model *K* is the *N*×*N* kinship matrix between individuals, *B* is the *P*×*P* matrix of genetic covariances between phenotypes and *E* is the *P*×*P* matrix of residual covariances between phenotypes. We estimate the covariance matrices *B* and *E* using a new C++ implementation of an EM algorithm^84^ included in the SBAT software (see **URLs**).

For the marginal heritabilities and genetic correlation analysis we used a realised relationship matrix (RRM) for the Kinship matrix (*K*). This RRM was calculated ^from^ the 8,428 nominally unrelated individuals using fastLMM (see **URLs**). We used the subset of imputed SNPs that were both assayed by the genotyping chips and included in the HRC reference panel, and so will essentially be hard-called genotypes. In addition, all SNPs with duplicate rsids were removed. PLINK (see **URLs**) was used for file conversion before input into fastLMM.

To estimate genetic correlations, we fit the model to several of the groupings of IDPs detailed in **Supplementary Table 12**. The estimated covariance matrices B and E were used to estimate the genetic correlation of pairs of IDPs. The genetic correlation between the ith and jth IDPs in a jointly analyzed group of IDPs is estimated as

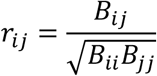

### Multi-trait association tests

We used a multi-trait mixed model to test each SNP for association with different groupings of traits detailed in **Supplementary Table 7**. The model has the form

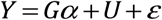

where *G* is an *N*×1 vector of SNP dosages and ^*α*^ is a 1×*P* vector of effect sizes. We fit the model using estimates of *B* and *E* from the “null” model with ^*α*=0^ and a leave one chromosome out (LOCO) approach for RRM calculation. We ran this test on the main set of 8,428 samples and on the replication samples. For the replication analysis we used the estimates of *B* and *E* from the main set of 8,428 samples. This test is implemented in the SBAT software (see **URLs**).

### Genetic association of IDPs

We used BGENIE v1.2 (see **URLs**) to carry out GWAS of imputed variants against each of the processed IDPs. This program was designed to carry out the large number of IDP GWAS required in this analysis. It avoids repeated reading of the genetic data file for each IDP and uses efficient linear algebra libraries and threading to achieve good performance. The program has already been used by several studies to analyze genetic data from the UK Biobank^85,86^. We fit an additive model of association at each variant, using expected genotype count (dosage) from the imputed genetic data. We ran associated tests on the main set of 8,428 samples and the replication samples.

### Identifying associated genetic loci

Most GWAS only analyze one or a few different phenotypes, and often uncover just a handful of associated genetic loci, which can be interrogated in detail. Due to the large number of associations uncovered in this study we developed an automated method to identify, distinguish and count individual associated loci from the 3,144 GWAS (one GWAS for each IDP). For each GWAS we first identified all variants with a -log10 p-value > 7.5. We applied an iterative process that starts by identifying the most strongly associated variant, storing it as a lead variant, and then removing it, and all variants within 0.25cM from the list of variants (equivalent to approximately 250kb in physical distance). The process was then repeated until the list of variants was empty. We applied this process to each GWAS using 2 different filters on MAF: (a) MAF > 0.1%, and (b) MAF > 1%. We grouped associated lead SNPs across phenotypes into clusters. This process first grouped SNPs within 0.25cM of each other, and this mostly produced sensible clusters, but some hand curation was used to merge or split clusters based on visual inspection of cluster plots and levels of LD between SNPs. For some clusters in **Table 1** we report coding SNPs that were found to be in high LD with the lead SNPs.

### Accounting for multiple IDPs

We adjusted the genome-wide significance threshold (-log10 p-value > 7. 5) by a Bonferroni factor (-log_10_(3144)=3.5) that accounts for the number of IDPs tested, giving a threshold of -log_10_ p > 11. This assumes (incorrectly) that the IDPs are independent and so is likely to be conservative, but we preferred to be cautious when analyzing so many IDPs.

### Genetic correlation analysis

We used LD score regression^87^ to estimate the genetic correlation between the IDPs studied in our analysis and 10 disease, personality or brain related traits. We gathered summary statistics for genome wide association studies of the neuroticism personality trait, autism spectrum and sleep duration and also 7 disease traits: attention deficit hyperactivity disorder, bipolar disorder, Alzheimer’s disease, major depressive disorder, schizophrenia, stroke and amyotrophic lateral sclerosis. The number of samples in each of these studies and the DOIs for the corresponding studies are provided in **Supplementary Table 13**.

For each IDP/trait pair, we used the LDSCORE regression software (v1.0.0) to compute the genetic correlation between the IDP and the trait, with linkage disequilibrium measurements taken from 1000 Genomes Project (provided by the maintainers of the LDSCORE regression software). We filtered the SNPs to include only those with imputation INFO >= 0.9 and MAF >= 0.1%. Only INFO scores for major depressive disorder, schizophrenia and attention deficit hyperactivity disorder were provided by the source studies, and so for these three analyses we applied the INFO threshold to both the SNPs from our study and also the source study. For the remaining 6 studies, an INFO filter was applied to the SNPs from our own study. Due to low levels of heritability of the functional edge IDPs, all of these were removed from this analysis. Since calculation of genetic correlation between traits only really makes sense if both traits are themselves heritable, we only used those IDPs with z-scores for significantly non-zero heritability greater than 4. In total we used 897 IDPs. To account for correlations between IDPs we used the raw phenotype correlation matrix to simulate z-scores (and associated tail probabilities) using samples from a multivariate normal distribution with that same correlation matrix.

### Analysis of enrichment of functional categories

We used the LDSCORE regression software to carry out the heritability enrichment partitioning analysis into different functional categories (see **URLs**). We used 24 functional categories:_coding, UTR, promoter, intron, histone marks H3K4me1, H3K4me3, H3K9ac5 and two versions of H3K27ac, open chromatin DNase I hypersensitivity Site (DHS) regions, combined chromHMM/ Segway predictions, regions conserved in mammals, super-enhancers and active enhancers from the FANTOM5 panel of samples. For each IDP, the enrichment of each functional category is summarized as the proportion of *h*^*2*^ explained by the category divided by the proportion of common variants in the category. For each IDP and each annotation we used the two-side enrichment p-value as reported by the LDSCORE regression software. We labeled those p-values as *enriched* or *depleted* depending on whether the enrichment estimate was greater or less than 1. We stratified these p-values accordingly into 23 groups of IDPs.

## Acknowledgements

The data used in this work was obtained from UK Biobank under Data Access Application 8107. We are grateful to UK Biobank for making the resource data available, and are extremely grateful to all UK Biobank study participants, who generously donated their time to make this resource possible. F.A-A acknowledges funding from the UK Medical Research Council and the Wellcome Trust via UK Biobank. KL.M.□and S.M.S. receive further support from the Wellcome Trust. JM acknowledges funding for this work from the European Research Council (ERC; grant 617306) and the Leverhulme Trust. GD acknowledges funding from the Medical Research Council UK (MR/K006673/1). We are grateful to: Bruce Fischl, Doug Greve and Matt Glasser for advice on FreeSurfer processing; Jon Diprose and Robert Esnouf for their advice and support with high performance computing; Stuart McRobert for help setting up and configuring the Oxford Brain Imaging Genetics browser; Tom Nichols and Anderson Winkler for discussions about the imaging confounds. For the genetic correlation analysis we used summary statistic data from several GWAS of brain related conditions as follows: the ISGC Cerebrovascular Disease Knowledge Portal, International Genomics of Alzheimer’s Project (IGAP), the Project MinE GWAS Consortium, the Social Science Genetic Association Consortium (SSGAC), the University of Exeter research group on Type 2 Diabetes, Obesity, Growth & Reproductive Ageing Genetics, the Psychiatric Genomics Consortium (PGC) and the ENIGMA consortium. We are grateful to these groups for making this data publicly available and to all the participants in these various studies.

## Author contributions

J.M and S.S conceived and supervised the work. F.A-A, K.M, G.D., S.S created the IDPs and confound covariates. L.E, K.S, S.Shi and J.M carried out the genetic association, heritability, genetic correlation and functional enrichment analysis and created the Oxford BIG browser. J.M, S.S, G.D, F.A-A, K.M, K.S and L.E interpreted the results and wrote the paper.

## URLs

Oxford BIG server http://big.stats.ox.ac.uk/

BGENIE https://jmarchini.org/bgenie/

SBAT https://jmarchini.org/sbat/

fastLMM https://github.com/MicrosoftGenomics/FaST-LMM PLINK http://www.cog-genomics.org/plink/2.0/

LDSCORE regression software https://github.com/bulik/ldsc PheWeb https://github.com/statgen/pheweb/

Various resources relating to the brain imaging in UK Biobank, including 3D-maps and connectome browsers for the group-ICA rfMRI analyses, and matlab code used to generate and apply the confound variables for this paper: http://www.fmrib.ox.ac.uk/ukbiobank/

UK Biobank showcase variables used for head positioning confounds and scan date:

http://biobank.ctsu.ox.ac.uk/showcase/field.cgi?id=25756

http://biobank.ctsu.ox.ac.uk/showcase/field.cgi?id=25757

http://biobank.ctsu.ox.ac.uk/showcase/field.cgi?id=25758

http://biobank.ctsu.ox.ac.uk/showcase/field.cgi?id=25759

https://biobank.ctsu.ox.ac.uk/showcase/field.cgi?id=53

Head bone density and mineral content measures:

*https://biobank.ctsu.ox.ac.uk/crystal/docs/DXAexplandoc.pdf*

GWAS summary statistics used for genetic correlation analysis

Major depressive disorder - https://www.med.unc.edu/pgc/

Schizophrenia - https://www.med.unc.edu/pgc/

Autism spectrum - https://www.med.unc.edu/pgc/

Attention deficit hyperactivity disorder and bipolar disorder - https://www.med.unc.edu/pgc/

Alzheimer’s disease - http://web.pasteur-lille.fr/en/recherche/u744/igap/igapdownload.php

Amyotrophic lateral sclerosis - http://databrowser.projectmine.com/

Stroke - PMC4818561 from http://cerebrovascularportal.org/informational/downloads

Neuroticism - https://www.thessgac.org/data

Sleep duration - http://www.t2diabetesgenes.org/data/

ENIGMA - http://enigma.ini.usc.edu/research/download-enigma-gwas-results/

